# Cycling DOF Factor mediated seasonal regulation of sexual reproduction and cold response is not conserved in *Physcomitrium patens*

**DOI:** 10.1101/2024.08.20.608156

**Authors:** Alexander G. Freidinger, Lauren A. Woodward, Jo Trang Bùi, Gillian Doty, Shawn Ruiz, Erika Conant, Karen A. Hicks

## Abstract

Many land plants have evolved such that the transition from vegetative to reproductive development is synchronized with environmental cues. Examples of reproduction in response to seasonal cues can be found in both vascular and nonvascular species; however, most of our understanding of the molecular events controlling this timing has been worked out in angiosperm model systems. While the organism-level mechanisms of sexual reproduction vary dramatically between vascular and nonvascular plants, phylogenetic and transcriptomic evidence suggest paralogs in nonvascular plants may have conserved function with their vascular counterparts (Holm et al. 2010; Zhao et al. 2019; Genau et al. 2021). Given that *Physcomitrium patens* undergoes sexual reproductive development in response to photoperiodic and cold temperature cues (Hohe et al. 2002), it is well-suited for studying evolutionarily conserved mechanisms of seasonal control of reproduction. Thus, we used publicly available microarray data to identify genes differentially expressed in response to temperature cues (Fernandez-Pozo et al. 2020). We identified two *CDF-like* (*CDL*) genes in the *P. patens* genome that are the most like the angiosperm *Arabidopsis thaliana* CDFs based on conservation of protein motifs and diurnal expression patterns. In angiosperms, DNA-One Finger Transcription Factors (DOFs) play an important role in regulating photoperiodic flowering, regulating physiological changes in response to seasonal temperature changes, and mediating the cold stress response (Imaizumi et al. 2005; Kloosterman et al. 2013; Fornara et al. 2015; Ridge et al. 2016; Ding et al. 2018; Blair et al. 2022). We created knockout mutations and tested their impact on sexual reproduction and response to cold stress. Unexpectedly, the timing of sexual reproduction in the *ppcdl* double mutants did not differ significantly from wild type, suggesting that the *PpCDLs* are not necessary for seasonal regulation of this developmental transition. We also found that there was no change in expression of downstream cold-regulated genes in response to cold stress and no change in freezing tolerance in the knockout mutant plants. Finally, we observed no interaction between PpCDLs and the partial homologs of FKF1, an *Arabidopsis thaliana* repressor of CDFs. This is different from what is observed in angiosperms (Fornara et al. 2009; Corrales et al. 2014; Song et al. 2016, Li et al. 2013; Han et al. 2015, Wang et al. 2023, Li et al. 2009), which suggests that the functions of CDF proteins in angiosperms are not conserved in *P. patens*.

## Introduction

The regulation of life sustaining processes in response to changing environmental conditions is fundamental for land plant survival and reproduction. Initiation of reproductive development in conjunction with favorable environmental conditions promotes fitness by increasing reproductive success (Garner and Allard 1920; Lang 1952). The molecular mechanisms driving seasonally regulated sexual reproduction in land plants are best understood in the model angiosperm, *Arabidopsis thaliana;* however, little is known about how these responses function in non-seed plant lineages. To this end, we use the model moss *Physcomitrium patens*, which shared a common ancestor with angiosperms ∼450 million years ago, to investigate the molecular mechanisms of reproduction in non-seed plants (Cove 2005). *P*. *patens*’ haploid dominant life cycle, ease of genetic manipulation, and the fact that little is known about reproduction in bryophytes make *P. patens* a rich model system to study evolutionary development of seasonal regulation of reproduction. Initiation of sexual development in *P. patens* occurs in the fall and can be induced in the laboratory under exposure to short days and cold temperatures (Hohe et al. 2002; Landberg et al. 2013; Hiss et al. 2017). Sexual reproduction begins with the development of sperm-bearing antheridia and egg-bearing archegonia, and fertilization leads to the formation of a sporophyte at the apex of a branch (Cove and Knight 1993).

There is remarkable conservation across land plants in terms of the transcriptional initiation of embryogenesis. Knotted-like homeobox (KNOX) transcription factors drive the haploid to diploid transition in all land plants and algae (Joo et al. 2018). This function is also conserved in *P. patens*, where *KNOX2* mutants undergo fertilization, but fail to undergo the typical sporophyte maturation, resulting in gametophyte tissue arising from diploid embryos with no meiosis (Sakakibara et al. 2013). In bryophytes, the haploid gametophyte is the dominant generation. Transition from haploid to diploid sporophyte tissue is mediated by the homeobox transcription factor BELL1, which initiates embryo formation and diploid sporophytes.

Overexpression of *BELL1* in *P. patens* results in formation of diploid embryos, without fertilization, while loss of *BELL1* results in sterility of eggs in archegonia (Horst et al. 2016). Expression of transcription factors required in the determination of male and female fates are epigenetically controlled in plants (Genau et al. 2021). Conserved epigenetic regulators, *HAG1* and *SWI3/B* mutants show reproductive defects in flowering plants and are critical for male and female gametangia development in *P. patens* (Genau et al. 2021).Taken together, a picture of conservation of genes required to initiate development of reproductive structures emerges across land plants. However, the degree of conservation of mechanisms driving the synchronization of reproductive development with seasonal and environmental cues is not understood across varied plant lineages, including bryophytes.

In contrast to *P. patens,* the molecular underpinnings of seasonal regulation of reproduction in *A*. *thaliana* are relatively well understood (Gendron and Staiger 2023). In *A. thaliana*, transcription of *FLOWERING LOCUS T (FT)* is promoted by the transcription factor CONSTANS (CO) in long days (Imaizumi and Kay 2006; Shim et al. 2017). CDF proteins repress the expression of *CO* leading to inhibition of sexual reproduction (Goralogia et al. 2017). However, CDF proteins are degraded late in the day by FLAVIN-BINDING, KELCH REPEAT, F-BOX 1 (FKF1), ZEITLUPE (ZTL), and LOV KELCH PROTEIN 2 (LKP2) in a GIGANTEA (GI) dependent manner (Sawa et al. 2007; Fornara et al. 2009). Additionally, *CDF* transcription is repressed late in the day by PRR5, PRR7, and PRR9, with *CDF* expression reaching the trough around dusk and returning to a high level at night (Nakamichi et al. 2007; Niwa et al. 2007; Fornara et al. 2009). During long days, there is increased repression of *CDF*, relieving the repression of *CO* and, subsequently, *FT* (Fornara et al. 2009).

A role for CDF-like genes as regulators of reproduction persists even in angiosperms distantly related to *A. thaliana*, though there are notable distinctions. For example, in day neutral *Solanum lycopersicum* (tomato), there are 5 CDF-like genes, *SlCdf1-5*. Of these, only overexpression of *SlCDF3* results in delayed flowering in both long day and short day conditions, suggesting a mechanism reminiscent of CDF regulation in long-day flowering plants (Xu et al. 2021). In both *Glycine max* (soybean), a short day responsive dicot, and *Oryza sativa* (rice), a short day responsive monocot, the GI-FKF1-CDF interaction appears to be conserved, yet the consequence of this interaction on flowering is reversed (Li et al. 2013; Han et al. 2015). While GI activates flowering in long day plants, the soybean *GI* homolog *E2* functions as an inhibitor of flowering (Wang et al. 2023). Likewise, while CDF1 delays flowering in *Arabidopsis*, overexpression of rice *DOF12* leads to early flowering in non-inductive long-day conditions (Li et al. 2009).

In addition to regulating sexual reproduction, CDFs are involved in the cold stress response in multiple angiosperms (Corrales et al. 2014; Fornara et al. 2015). AtCDF3 regulates the expression of several genes involved in the response to extreme temperatures, drought and osmotic stress, including several central abiotic stress regulators like *Late Embryogenesis Abundant* (*LEA*), *Cold Binding Factors* (*CBF*), *Drought Response Element Binding Factors* (*DREB*) and *Salt Tolerance Zinc Fingers* (*ZAT*) (Corrales et al. 2017). This mechanism utilizes both AtCDF3-GI dependent and AtCDF3-GI independent pathways, reflecting an integration between the seasonal regulation of sexual reproduction and cold stress response regulation. Interestingly, *LEA* expression is also increased in *P. patens* exposed to cold temperatures, suggesting that *PpDOF* levels may drive cold-stress dependent increases in LEA expression in *P patens*, as well (Minami et al. 2005; Takezawa et al. 2015).

There are three domains of interest within the *Arabidopsis* CDF proteins: the DOF domain, the TOPLESS binding domain (Goralogia et al. 2017), and the GI/FKF binding domain (Imaizumi et al. 2005; Sawa et al. 2007). The characteristic DOF domain of AtCDF1 binds to the repeated-AAAG-DNA motif within the *FT* and *CO* promoters (Imaizumi et al. 2005), while the TOPLESS binding domain allows AtCDF1 to interact with TOPLESS, thereby repressing transcription of *FT* and *CO* (Goralogia et al. 2017). Together, AtGI and AtFKF1 bind to and initiate degradation of the AtCDF proteins via the ubiquitin ligase activity of AtFKF1 in reproductively inductive conditions or conditions with significant cold stress (Nelson et al. 2000; Lee et al. 2018; Lee et al. 2019). The N-terminus of GI binds the PAS/LOV domain of AtFKF1. Subsequently, six Kelch repeats of AtFKF1 interact with the AtCDFs (Sawa et al. 2007; Lee et al. 2018).

While many genes in the photoperiodic flowering pathway are conserved between *A. thaliana* and *P. patens* (including *CDF, CO, ELF3, ELF4*, and others), *GI* is conspicuously absent in *P. patens* (Shimizu et al. 2004; Zobell et al. 2005; Okada et al. 2009; Holm et al. 2010). Also absent in *P. patens* is a complete *FKF1* homolog. While there are several proteins that contain the domains found within FKF1, no single protein contains a PAS/LOV, F-box, and Kelch repeat domain (Holm et al. 2010). Taken together, these observations suggest an alternative means of regulating CDF activity and perhaps an alternative mechanism for regulating sexual reproduction in response to seasonal cues in *P. patens*.

In this study, we describe the lack of conservation of CDF regulation and function between *P. patens* and angiosperms. First, *P. patens Cycling DOF Factor like* (*PpCDL)* genes were identified using multiple phylogenetic analyses. Several of these genes are upregulated both in cold temperatures and in reproductive gametophores in publicly available microarray datasets (Hiss et al. 2014; Fernandez-Pozo et al. 2020). Further, diurnal and circadian-regulated expression patterns of one *PpCDL* gene, *PpCDL1* was similar to *AtCDF*s. These observations hinted at a potential conservation of function between *PpCDL1* and *AtCDF*s. To test this, *ppcdl* double knockout mutants were generated using the CRISPR-Cas9 genome editing system. The *ppcdl* double knockouts showed no change in reproductive timing when compared to wild type plants, indicating that either these two PpCDLs do not appear to play a major role in sexual reproduction or that PpCDLs act alongside redundant pathways to control seasonal regulation of sexual reproduction. Additionally, cold responsive genes were upregulated after exposure to near-freezing temperatures in the *ppcdl* double knockouts, similar to wild type. Furthermore, although PpCDL1 contains a conserved GI/FKF binding domain, we did not observe interaction with FKF-like proteins in a yeast two-hybrid assay. Together, our results indicate that CDF-like genes in *P. patens* are not functionally conserved relative to angiosperms.

## Results

### Identification of *P. patens CDF-like* (*CDL*) genes

To identify *P. patens* genes that are differentially expressed in reproductively inductive conditions, we examined publicly available microarray data and found that several annotated CDF-like homologs were expressed at higher levels in response to cold stress and in plants at more sexually mature stages of development (Hiss et al. 2014; Beike et al. 2015; Fernandez-Pozo et al. 2020). Multiple angiosperm CDFs are known to be involved in cold stress response in addition to sexual reproduction. The PpCDLs were expressed at higher levels in the 24 hours of exposure to cold (0 °C) (Figure 1a). Additionally, these PpCDLs were upregulated in adult gametophores compared to juvenile gametophores (Figure 1b). These two findings suggest that the PpCDLs may be involved in both initiation of sexual reproduction and response to cold stress, similar to *Arabidopsis* CDFs.

**Figure 1.**
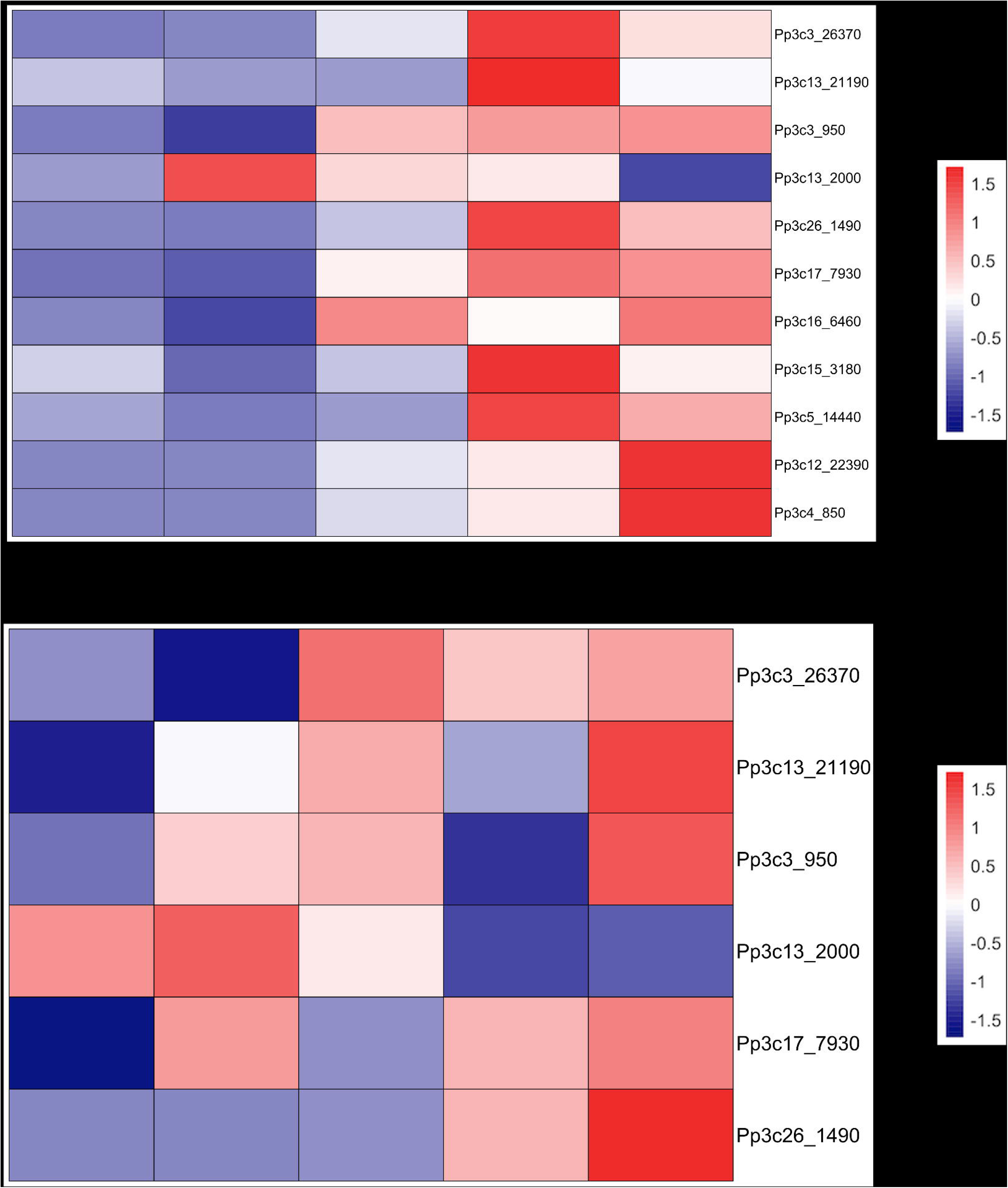
CDF-like homologs are expressed at higher levels in response to cold stress and in sexually mature gametophores. A. *PpDOF* expression increases in response to cold temperature (0 °C), with peak expression varying between genes. *Pp3c13_2000* expression peaked earliest and quickly returned to baseline. Expression of *Pp3c13_21190, Pp3c15_3180, Pp3c3_26370, Pp3c26_1490,* and *Pp3c5_14440* peaked 8 hours after exposure to cold temperatures and decreased at 24 hours. *Pp3c12_22390* and *Pp3c4_850* expression was highest 24 hours after cold exposure. Data from Fernandez-Pozo et al. (2020). B. *PpDOF* expression increases through sexual development. Peak expression varied between *PpDOFs,* with expression typically highest in gametangia containing adult gametophores. Data presented from Reute accession obtained from Perroud et al. (2018).

To determine the relationships between the 6 AtCDF proteins and PpDOF proteins that were upregulated in response to cold temperatures, we obtained homologous CDF peptide sequences from multiple angiosperm species, two green algae, two mosses, and a liverwort to reconstruct phylogenetic relationships (Figure 2A and Supplemental Figure 1). The tree revealed six *P. patens* proteins (Clade I) that are more closely related to the AtCDFs (posterior probabilities between 78.8% and 99.9%) than the other 10 *P. patens* proteins (Clade II). Next, we used MEME (Bailey et al. 2015) to identify domains in the *P. patens* and *A. thaliana* proteins (Figure 2B). All six *P. patens* proteins within Clade I contain the common CX2CX21CX2C zinc finger domain structure present in AtCDFs. In addition to the DOF domain, an N-terminal TOPLESS-binding domain was detected in four proteins, while a conserved C-terminal GI/FKF1 binding domain (Imaizumi et al. 2005; Kloosterman et al. 2013; Ridge et al. 2016) was found in the two proteins most closely related to *Arabidopsis* CDFs. Both the TOPLESS-binding and GI/FKF1-binding domains are present in *Arabidopsis* CDFs; however, these two domains were not found within the same protein in *P. patens*. While *TOPLESS*-like genes are present in the *P. patens* genome (Goralogia et al. 2017), neither a *GI* homolog nor a complete, tripartite *FKF1* homolog are found in the *P. patens* genome; however partial *FKF1* homologs are present (Holm et al. 2010, Figure 2B).

**Figure 2.**
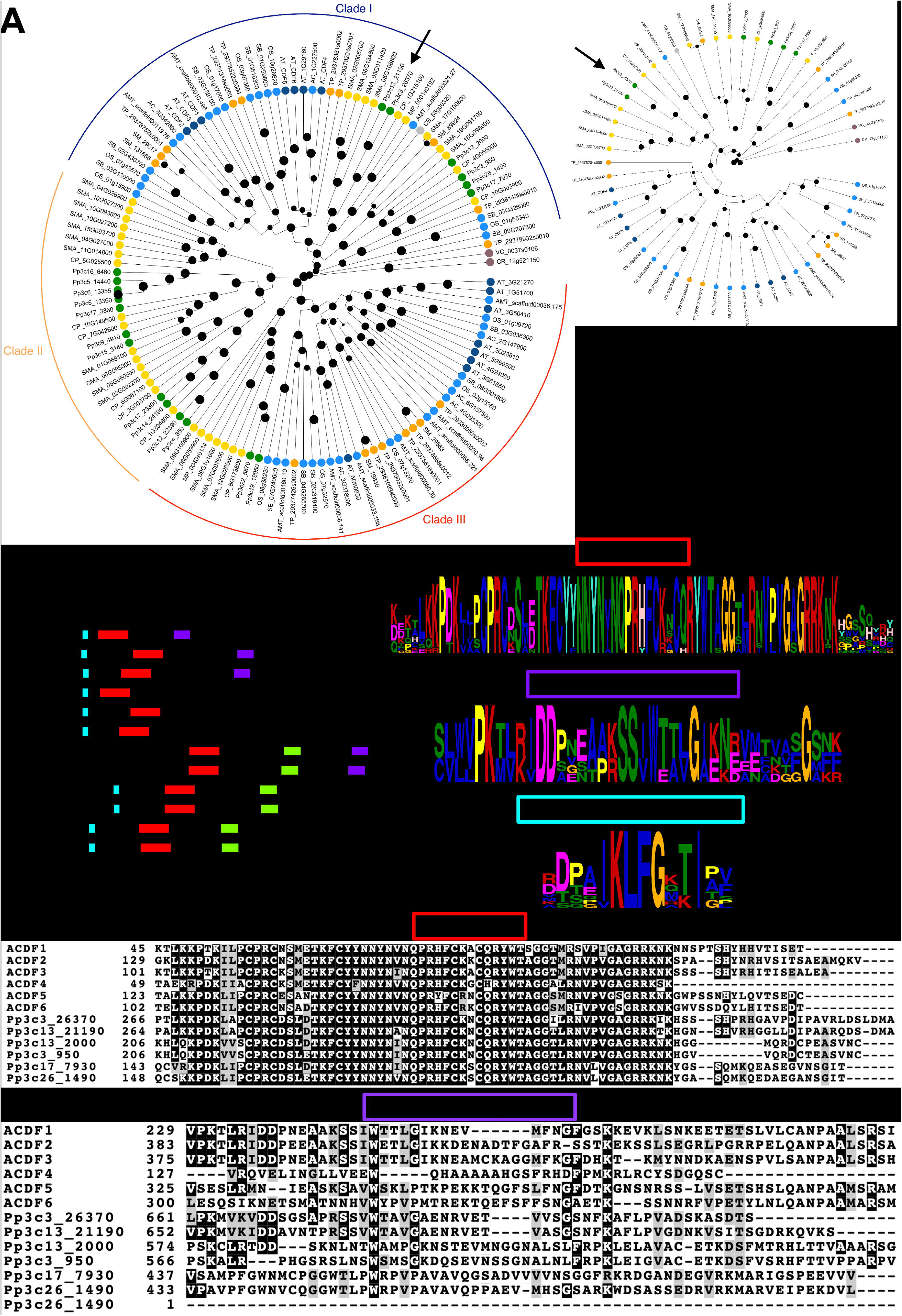
Phylogenetic analysis of PpDOFs reveals two PpCDL genes. A. Bayesian phylogenetic tree highlighting relationships between AtCDFs and other plants’ CDF homologs. *P. patens* genes of interest are indicated by the arrow. *A. thaliana-* blue. *P. patens* - green. *S. fallax* and *M. polymorpha-* yellow. Algae - gray. Other angiosperms - pale blue. Nodes are sized according to posterior probability, with larger dots representing higher posterior probability. Tree generated using BEASTv1.10.4. B. Domains found in AtCDFs, PpCDLs, and PpDOFs. Red box - DOF domain. Light blue box - TOPLESS domain. Purple box - FKF1 - GI binding domain. Consensus logos for three conserved motifs: DOF domain, FKF1-GI binding domain, and TOPLESS binding domain generated from MEME analysis. GI binding domain has been identified in *A. thaliana* (Sawa et al. 2007), and the TOPLESS binding domain has characteristic IKLFG sequence (Goralogia et al. 2017). C. DOF and FKF1-GI binding domain amino acid alignment show these domains to be conserved between the AtCDF and PpCDL proteins.

### *PpCDL1* is diurnally expressed

In angiosperms, CDFs are expressed diurnally (Fornara et al. 2009; Corrales et al. 2014). In order to determine which, if any, of the *P. patens* DOFs might also be diurnally expressed, we assessed expression of Clade I and Clade II *P. patens* transcript levels over a 24-hour period. While *Pp3c3_26370* was expressed diurnally, *Pp3c13_21190* did not exhibit a diurnal rhythm. This result was consistent across multiple growth conditions (Figure 3A, Supplemental Figure 1A). In fact, except for *Pp3c3_26370*, none of the *P. patens DOF* genes that we tested were expressed diurnally (Figure 3A). Given that *Pp3c3_26370* exhibited diurnal expression, and that both *Pp3c3_26370* and its closest homolog *Pp3c13_21190* were found to contain a conserved C-terminal GI/FKF1 binding domain, we concluded that *Pp3c3_26370* is the *P. patens* gene most closely related to AtCDFs and termed *Pp3c3_26370* and *Pp3c13_21190* as *Cycling DOF Factor-like (CDL) 1* and *2*, respectively.

**Figure 3.**
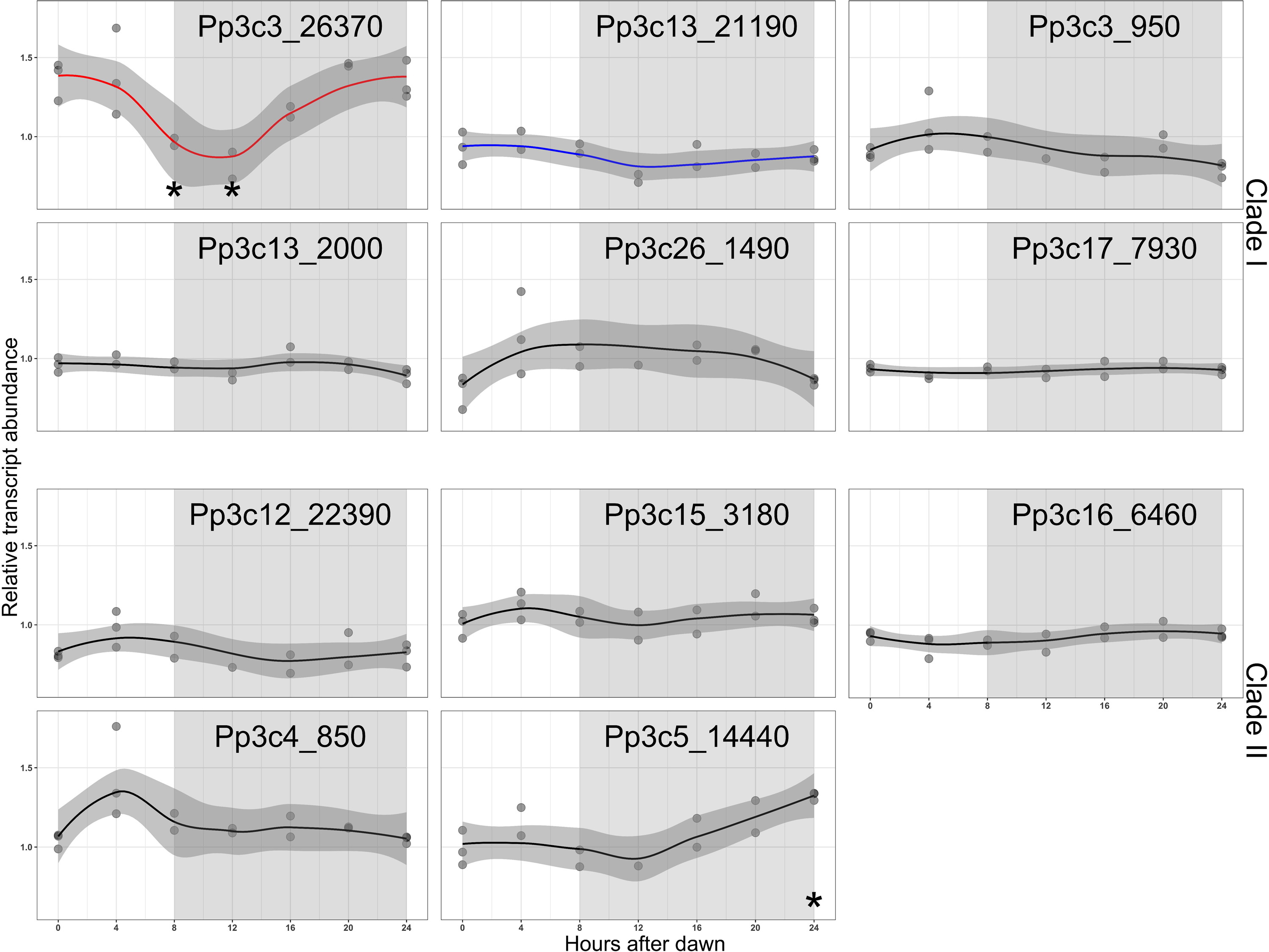
*PpCDL-1* is expressed diurnally. *HIBCH* normalized qRT-PCR expression curves for 11 *PpDOF* genes in conditions that induce gametangia formation (8L16D 15°C). Only Pp3c3_26370 showed strong diurnal expression. Quantitative PCR was carried out using SYBR Green MasterMix and data was analyzed using the relative standard curve method. All graphs plotted with the same y-axis. Asterisks represent time points that are significantly different from time 0 (p≤0.05) with both ANOVA and Tukey HSD tests.

### Identification of putative PpCDL regulators through phylogenetic analysis

In *A. thaliana*, CDF protein degradation is mediated through interaction with the FKF1 Kelch repeats and the GI PAS domain (Lee et al. 2018; Lee et al. 2019). Conservation of the GI/FKF1-binding domain in the two PpCDLs suggested that this protein degradation pathway might also be conserved. A BLAST search identified no proteins resembling GI in the *P. patens* proteome, consistent with the findings of Holm et al. (2010). However, we identified 15 proteins in *P. patens* with sequence similarity to AtFKF1 and constructed phylogenetic relationships using a Bayesian tree to determine the *P. patens* proteins most closely related to AtFKF1 (Figure 4A). From this analysis, we identified four proteins that are closely related to AtFKF1 and its sister proteins LKP2 and ZTL. Two of these proteins contain two PAS/LOV domains, which are involved in light and temperature signaling (Holm et al. 2010). The other two closely related proteins have an F-box and 5 Kelch repeats, which may be involved in facilitating degradation of other proteins. While Holm et al. (2010) also identified two proteins with F-box and Kelch domains, they found only two Kelch repeats in their analysis, perhaps due to inaccuracies in earlier versions of the *P. patens* genome annotation.

**Figure 4.**
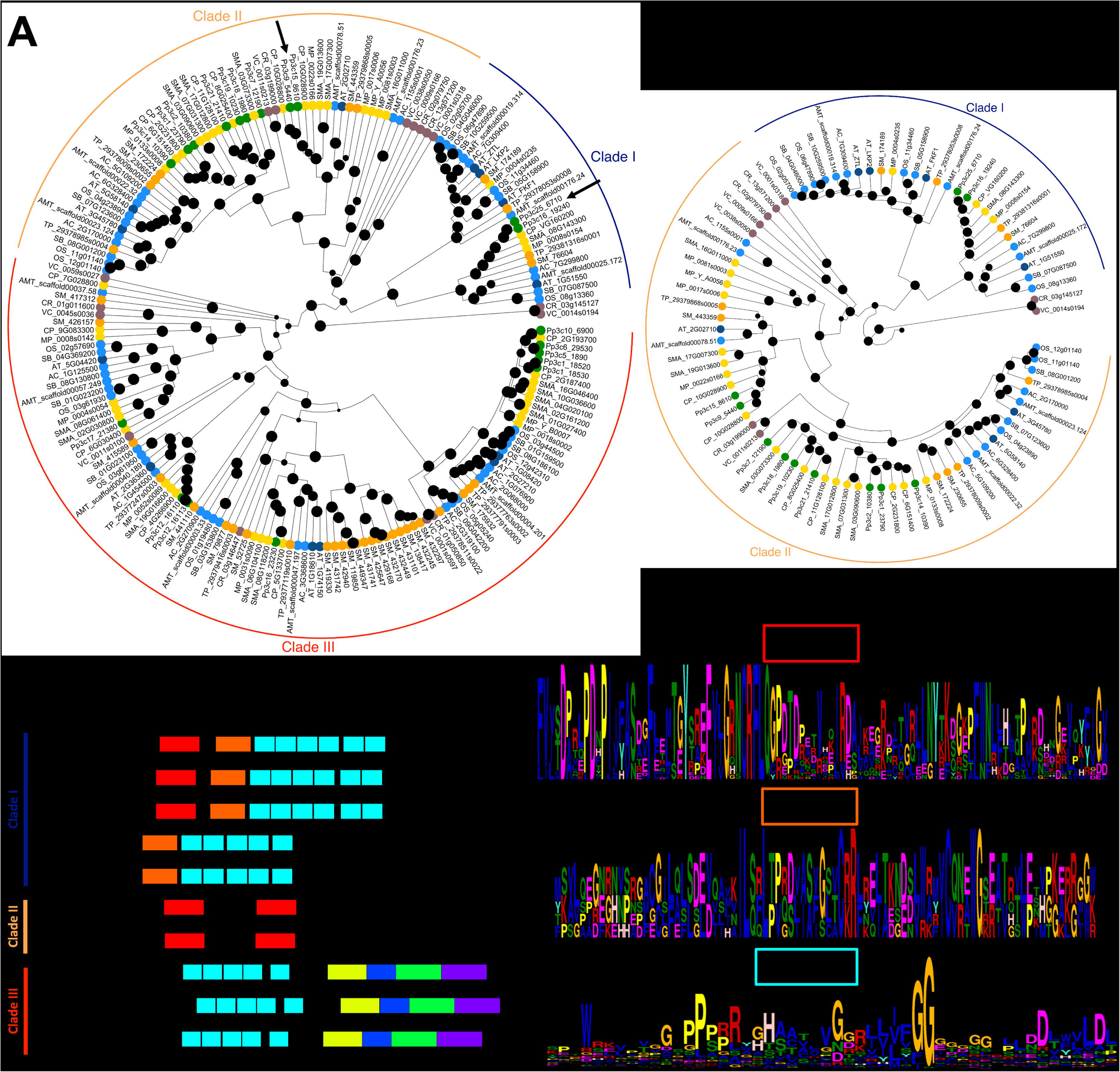
Phylogenetic analysis reveals four partial homologs of FKF1 in *P. patens*. A. Bayesian phylogenetic tree highlighting relationships between AtFKF1, AtLKP1 (AT2G18915), AtZTL (AT5G57360) and other plants’ AtFKF1 homologs. *P. patens* genes of interest are indicated by the arrows. *A. thaliana* – dark blue. *P. patens* - green. *S. fallax* and *M. polymorpha -* yellow. Algae - gray. Other angiosperms - light blue. Dots on nodes are sized according to the posterior probability of the node with the larger dots representing a higher posterior probability (posterior probabilities 0.5109-1). Tree generated using BEASTv1.10.4. B. Domains found in AtFKF1, AtLKP2, AtZTL, PpFKLs, and PpPASPAS. Red box - PAS-LOV domain. Orange box - F-box domain. Light blue box - FKF1-Kelch repeat domains.

### PpCDLs dimerize but do not interact with FKF1 partial homologs

To test the interaction between the FKF-binding class of PpCDLs and components of the *P. patens* FKF1 partial homologs (Fig. 4B), we performed a yeast two-hybrid assay with PpCDL genes, a twin-LOV gene, and two F-box/Kelch genes. No interaction was detected between PpCDL and any of the *P. patens* FKF1 partial homologs (Fig. 5A). Because the F-box degrades CDFs within the FKF1 complex in *A. thaliana*, it is possible that interaction between PpCDLs and F-box/Kelch proteins would not be detected in yeast two-hybrid experiments if the F-box were to degrade the CDL before reporter gene expression. Therefore, we tested interactions between FKF1-binding PpCDL1, a twin-LOV protein, and the Kelch repeats of *P. patens* lacking an F-box. None of these components interacted with one another in yeast (Fig. 5B), suggesting that the role of *A. thaliana* FKF-1 in degrading CDFs as an upstream regulator of sexual reproduction is not shared with *P. patens* FKF1 partial homologs.

**Figure 5.**
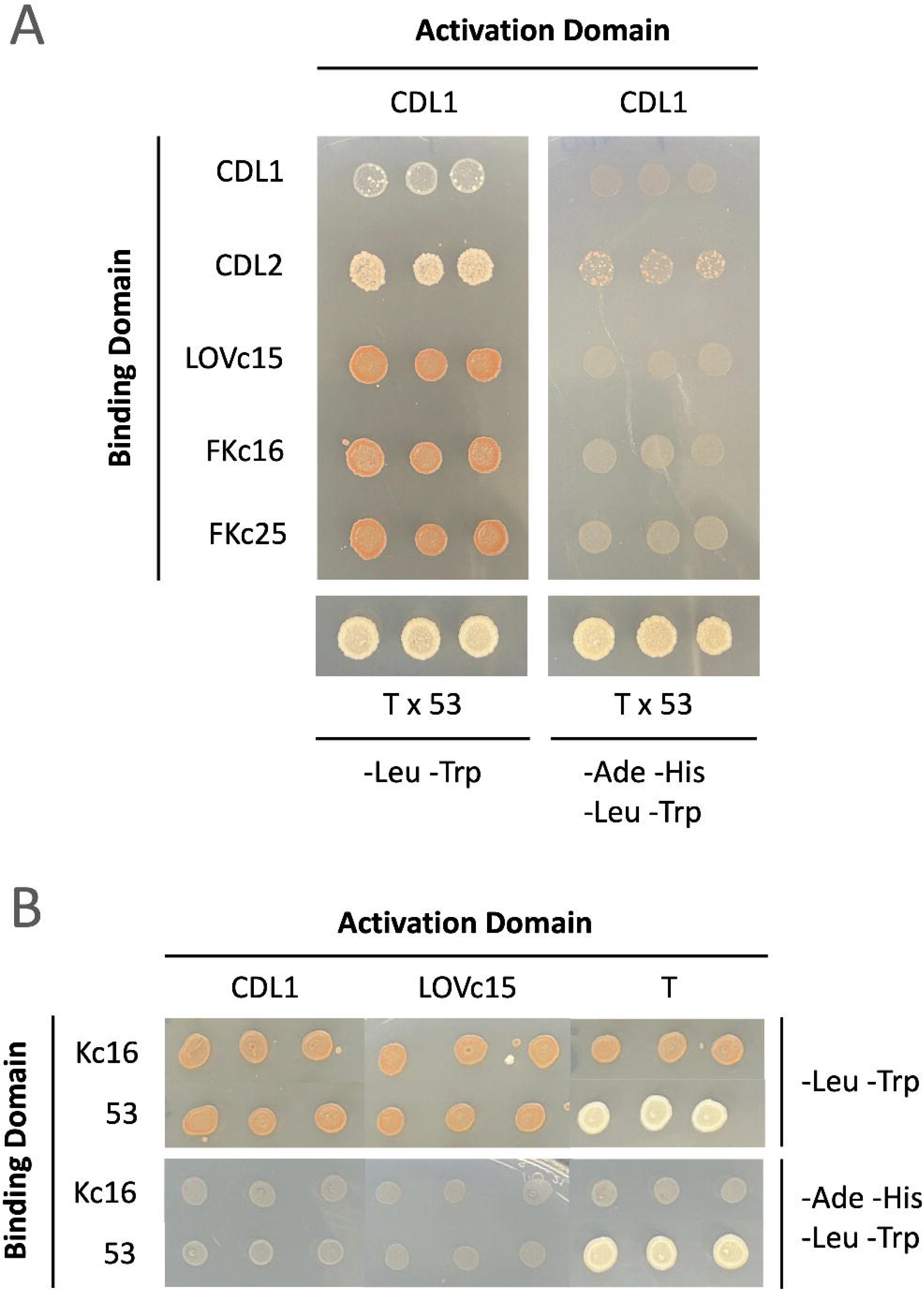
PpCDL protein interactions via yeast two-hybrid assay. A. *PpCDL1* and *PpCDL2* interact in yeast, suggesting the formation of a heterodimer *in vivo*. B. *PpCDLs* do not interact with *FKF1* partial homologs, even in the absence of an F-box. T x 53 is the positive control. - Leu/-Trp plates select for diploid yeast; -Ade/-His/-Leu/-Trp plates select for protein interaction.

Interestingly, the PpCDL1 and PpCDL2 formed a heterodimer in yeast (Fig. 5B, Fig. S1). This dimerization may play a role in the regulation or function of PpCDLs, consistent with reported examples of DOF protein dimerization in *A. thaliana*, wheat, and corn (Yanagisawa 1997; Sani et al. 2018; Merlino et al. 2023).

### PpCDLs are not required for sexual reproduction in *P. patens*

To discern if the *PpCDL* genes are required for the seasonal regulation of sexual reproduction in *P. patens*, *PpCDL1* and *PpCDL2* mutant strains were generated utilizing CRISPR-Cas 9 gene editing (Lopez-Obando et al. 2016). Knockout strains were successfully identified in the Cholsey-K5 accession by Sanger sequencing (Supplemental Table 6). Compared to other accessions, Cholsey-K5 was chosen because of its consistency in initiating sexual reproduction only in response to seasonal cues (unpublished). Plants were grown in long day (16L8D) conditions at 22°C until all plants were vegetatively mature. Then, plants were transferred to 12L12D 22°C conditions to assess for the differences in reproductive development between mutants and wild type, as this intermediate day length condition would allow us to observe either late or early reproduction in the knockout mutants. However, the *ppCDL1/2* double mutants exhibited little to no difference in reproductive timing compared to the WT (Figure 6). This suggests that the *PpCDLs* are not required for proper regulation of sexual reproduction in *P. patens*.

**Figure 6.**
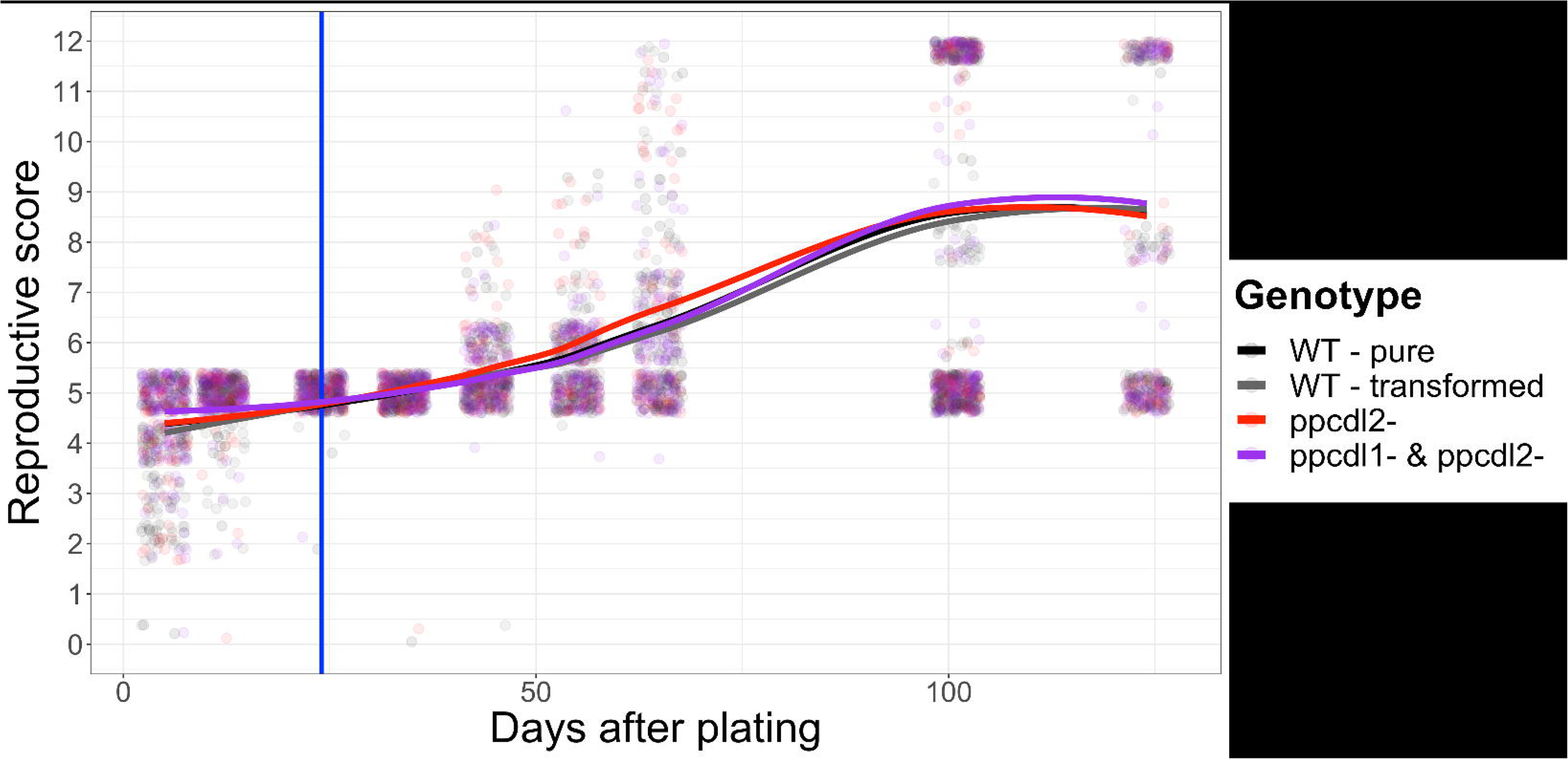
*ppcdl* knockout mutants reproduce similar to wild type. Reproductive timing was scored using a qualitative developmental stage from spores to mature brown sporophytes (Supplemental Table 5). Plants were germinated and grown in 16L8D 22°C before shift to 12L12D 22°C (denoted with vertical blue line). No significant differences between genotypes were observed between wild type Cholsey-K5, transformed Cholsey-K5 without mutations, *ppcdl2-*, and *ppcdl1-ppcdl2-* double mutants (ANOVA, p>0.05).

### *PpCDLs* are not required for cold stress response in *P. patens*

As angiosperm CDFs have been shown to be involved in the response to cold stress (Fornara et al. 2015; Corrales et al. 2017), we assessed the levels of expression in downstream genes involved in the *P. patens* response to cold stress in wild type and *ppCDL1/2* backgrounds (Sun et al. 2007). We observed strong induction of cold responsive genes after 3 days at 0°C in both wild type and *cdl* knockout strains (Figure 7), indicating that the *PpCDLs* are not required for proper cold responsive gene expression.

**Figure 7.**
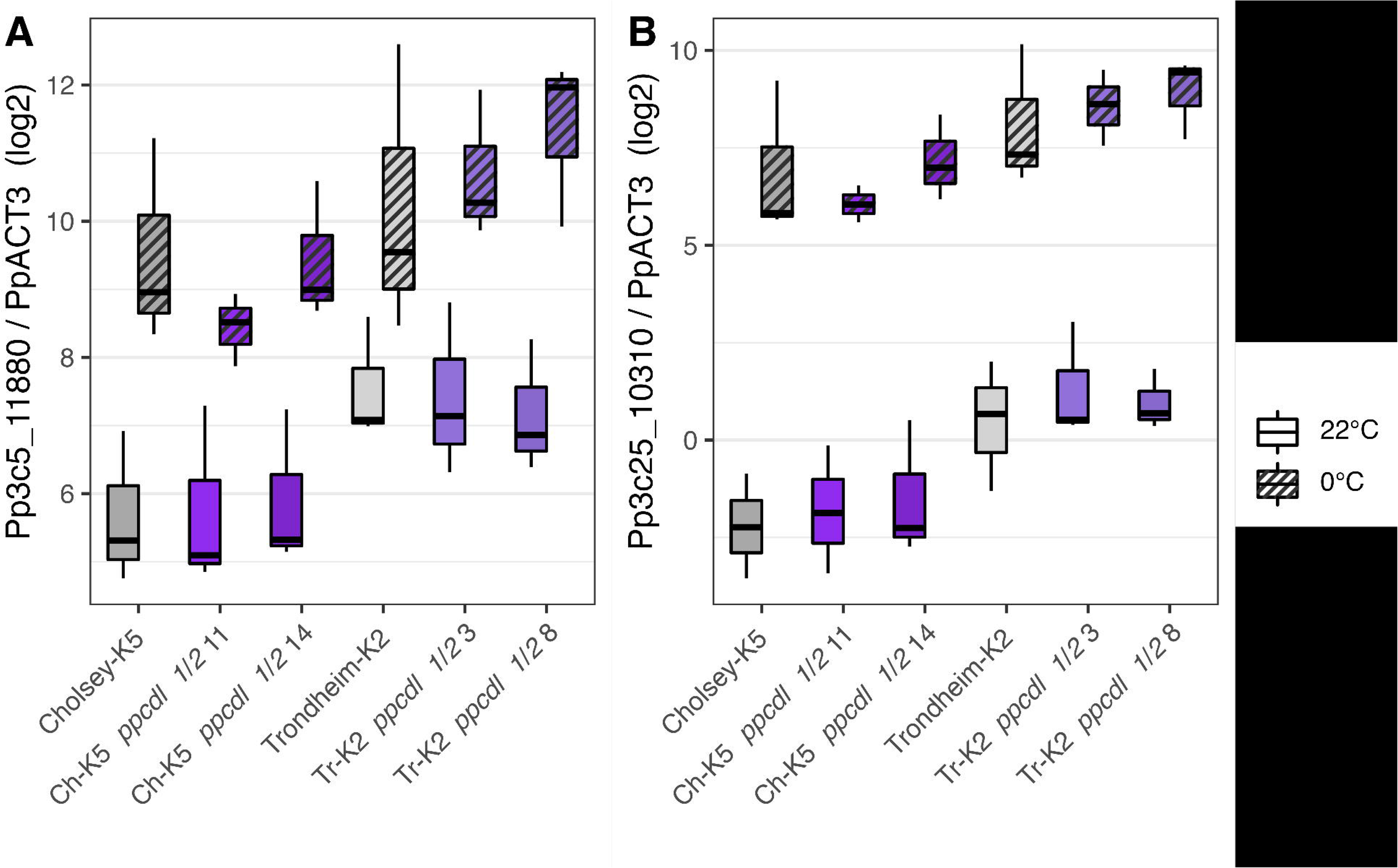
Expression of cold-responsive genes did not different between wild type and *ppcdl* double knockout mutants. *ACTIN3* normalized qRT-PCR expression curves for two cold responsive genes, Pp3c5_11880 and Pp3c25_10310 using PowerUP SYBR Green MasterMix and analyzed via the ΔΔCt method. Plants were grown in 16L8D 22°C for one month prior to shift to 0°C for 74 hours, with tissue collected 10 hours after dawn. Expression was evaluated in two *ppcdl* double knockout mutants in each of two *P. patens* accessions.

## Discussion

CDF homologs occupy some role in photoperiodic regulation of flowering across angiosperms (Corrales et al. 2014; Fornara et al. 2015; Song et al. 2016). Even in the alga *Chlamydomonas reinhardtii,* the DOF-CO module integrates environmental cues to regulate synchronous progression through the cell cycle in response to day length (Lucas-Reina et al. 2015). Of course, CDF function is most well understood in *A. thaliana*, where CDF represses flowering via inhibition of *CO* transcription. In tomatoes, *SlCDF3* also represses flowering (Xu et al. 2021). However, in short day responsive plants, while CDF is conserved and participates in flowering regulation, the consequences of CDF activity do not appear to be conserved (Li et al. 2009; Wang et al. 2023). The discrepancies among distantly related plants within an otherwise conserved module suggest an incomplete understanding of an interesting evolutionary history. To develop a more complete picture of the evolutionary development of seasonal reproduction in land plants, this study investigated the conservation of CDF function in a nonvascular model system, *P. patens*.

Through phylogenetic analysis, we identified a group of PpDOFs and narrowed this group down to two of these genes of interest: *PpCDLs.* We proceeded to generate *ppcdl1*, *ppcdl2,* and *ppcdl1/ppcdl2* mutants, and we determined that the *ppcdl* mutants do not exhibit a reproductive phenotype, suggesting that the *PpCDLs* are not involved in sexual reproduction in *P. patens*. Previous knockouts of *PpDOF1* (Pp3c17_7930) and *PpDOF2* (Pp3c26_1490) genes in Gransden result in a 2-fold reduction of protonemal branching in early stages of vegetative growth following growth from individual protoplasts (Sugiyama et al. 2012). We did not observe a similar phenotype in our *ppcdl1/2* (Pp3c3_26370 and Pp3c13_211990) knockout plants when grown either from macerated tissue (Figure 6) or directly from spores (data not included). Further, our analyses did not reveal any significant delay in vegetative development prior to the initiation of sexual reproduction.

Additionally, we found that *PpCDL* regulation is not conserved between angiosperms and *P. patens*. AtCDFs are regulated by FKF1 via binding of the Kelch repeat domain and degradation by the F-box domain (Imaizumi et al. 2005). In contrast, yeast two-hybrid analyses do not show interaction between PpCDL1 and *P. patens* Kelch repeats (Figure 5), suggesting that the regulation of CDFs via FKF1 is not conserved in *P. patens*. Taken together, this suggests PpCDLs and angiosperm CDFs likely target different genes and may be degraded via different mechanisms.

Interestingly, PpCDL1 and PpCDL2 formed a heterodimer in the yeast two-hybrid analyses. This is consistent with the studies of maize DOF proteins, where Yanagisawa (1997) demonstrated DNA-dependent DOF protein heterodimerization. Similar results were observed in *in vitro* gel shift assays with *A. thaliana* DOF-domains (Sani et al. 2018). More recently, DOF dimerization was also observed in sweet potato using yeast two-hybrid analyses (Zhang et al. 2023). How dimerization of DOF proteins – including PpCDL – contributes to physiological responses *in vivo* is not yet understood. It is likely that this broader model of regulation of DOF transcription factor activity through heterodimerization is conserved between *P. patens* and other land plants. However, the functional consequence of this dimerization requires further investigation. In any case, our results do not suggest that PpCDL dimerization is relevant to regulation of seasonal reproduction or to the cold stress response.

Functional studies of AtCDF3 by Corrales et al. (2017) and Fornara et al. (2015) showed AtCDF’s role in response to various abiotic stress responses, including cold stress. The molecular mechanisms of cold response in *P. patens* are similar to *A. thaliana* with some distinct differences in regard to CBF and DREB1 functions (Sun et al. 2007). We placed our *ppcdl1/ppcdl2* mutants in cold stressed environments and found that the expression of *LEA* genes was similarly increased in both wild type and knockout mutant plants, suggesting that *PpCDLs* are not required for response to cold stress.

In sum, established functions of CDFs in other land plants are not recapitulated in this work. This was unexpected, as the *PpCDLs* share sequence similarity with angiosperm CDFs and are expressed in a similar diurnal pattern as angiosperm CDFs. One explanation for the seeming lack of functional conservation between PpCDLs and CDFs of other land plants could be functional redundancy with one or several of the remaining four PpDOFs (Figure 2A). These four homologs contain a TOPLESS binding domain, but no FKF1/GI binding domain (Figure 2B). It is possible that these other PpDOFs compensate in the *ppdcl1/2* plants to maintain typical patterns of seasonal reproduction as well as the cold stress response. However, this would suggest that a FKF1/GI binding domain is irrelevant to seasonal regulation of reproduction in *P. patens.* Thus, there is unlikely to be functional conservation of PpCDL regulation and, by extension, conservation in the mechanisms of regulation of seasonal reproduction in *P. patens*. This is perhaps not surprising, given the loss of GI and the absence of a complete FKF1 homolog containing all three domains.

Another possibility is that the GI/FKF1-CDF-CO regulatory nexus has been lost in the moss lineage. The bryophyte lineage is monophyletic, and includes mosses, liverworts, and hornworts (Su et al. 2021). FKF1 and GI homologs are present in both the hornwort genus *Anthoceros* and in the liverwort *Marchantia polymorpha* (Linde et al. 2017, Laosuntisuk et al. 2023). However, Linde et al. (2017) found that GI and FKF are only present in one deeply divergent species of moss, *Takakia lepidozioides,* out of the 41 species of moss that were studied, suggesting that the FKF1-GI regulatory module was lost early in the moss lineage. In the liverwort *M. polymorpha*, GI and FKF1 are conserved and interact to regulate photoperiodic growth phase transitions, including the vegetative to reproductive transition (Kubota et al. 2014). Knockout of *MpGI* or *MpFKF* both abolished the development of gametangia, while overexpression caused a transition to gametangia development even under conditions where wild-type plants do not typically form gametangia. *M. polymorpha* also has a homolog of CDF which has the domains described in the *AtCDF* gene, including the FKF1 and GI-interaction domains and the TPL binding site (Kubota et al. 2014); However, the function of *MpCDL* in reproduction has not been investigated (Goralogia et al. 2017). It seems that the GI-FKF1 regulatory system precedes the transition to land and was co-opted by early land plants for regulating timing of reproduction but was lost in the moss lineage (Kubota et al. 2014; Goralogia et al. 2017). Consistent with this study, loss of GI and FKF1 in mosses may have allowed degeneration of CDF regulation and its role in regulating reproductive timing. Future work in this area will undoubtedly focus on identifying what alternative system *P. patens* has evolved to address gametangia development in response to daylength and temperature cues.

## Methods

### Amino acid sequence retrieval

AtCDF1-6 amino acid sequences were compiled using TAIR database (arabidopsis.org) via keyword searches and used to BLASTP search for homologs in Phytozome v12 (Holm et al. 2010; Zhao et al. 2019; Genau et al. 2021). Primary protein sequences were selected for *Selaginella moellendorffii* (v1.0), *Arabidopsis thaliana* (TAIR10), *Gossypium raimondii* (v2.1), *Populus trichocarpa* (v3.0), *Glycine max* (Wm82.a2.v1), *Amborella trichopoda* (v1.0), *Sphagnum fallax* (v0.5), *Physcomitrium patens* (v3.3), *Marchantia polymorpha* (v3.1), *Aquilegia coerulea* (v3.1), *Oryza sativa* (v7_JGI), *Solanum lycopersicum* (iTAG2.4), *Sorghum bicolor* (v3.1.1), *Chlamydomonas reinhardtii* (v5.5), and *Volvox carteri* (v2.1). AtCDF1-6 amino acid sequences were also used to search for homologs in *Chara braunii* (v1.0) and *Anthoceros* (taxid: 3233) species using BLASTP on NCBI (https://blast.ncbi.nlm.nih.gov/Blast.cgi). Sequence names were edited to only include family and species name abbreviation (Supplemental Table 1) and location of sequence in the genome.

### Phylogenetic analysis

Sequences were aligned using MUSCLE (https://www.ebi.ac.uk/Tools/msa/muscle/) and alignment was visualized using AliView (v1.25, https://ormbunkar.se/aliview/) to ensure that no large segments of poor alignment were present (Supplemental Figure 2A). Improved posterior probabilities in resulting Bayesian trees support elimination of sequences with limited homology (Supplemental Figure 2B). Aligned sequences were converted to BEAST input format using BEAUti (v1.10.4, https://beast.community), with default settings used except for the following: Tree Prior: “Speciation: Yule Process” and MCMC Length of chain: “30000000”. Bayesian trees were constructed using BEAST (v1.10.4). TreeAnotator (v1.10.4) was used to compile trees with default settings and the compiled tree file was analyzed and annotated in R Studio using ggtree (v2.4.1), phytools (v0.7), and treeio (v1.14.3) packages. The same protocol was used for phylogenetic analysis of FKF1/ZTL/LKP2 related amino acid sequences.

### Protein motif analysis

*P. patens* and *A. thaliana* amino acid sequences were uploaded to MEME (http://meme-suite.org/tools/meme) to identify conserved protein motifs (Bailey et al. 2015). MEME was used in Classic Mode with default settings except for the following: Site Distribution: “Any Number of Repetitions (anr)”, Number of motifs: “4”, minimum motif width: “6”, and maximum motif width: “100”. Consensus logos, motif locations, and statistical output were saved. Domains were identified using InterProScan (https://www.ebi.ac.uk/interpro/search/sequence/) and previously published literature.

### Plant material and growth conditions

*Physcomitrium patens* was grown on solid BCD medium (0.8% agar, 1 mM MgSO4, 1.85 mM KH2PO4, 10 mM KNO3, 45 μM FeSO4, 1 mM CaCl2, 1× Hoagland’s Number 2 solution) with 16 hours of light and 8 hours dark at 22°C unless otherwise noted.

### RNA extraction and cDNA synthesis

Tissue samples were frozen in liquid nitrogen and mechanically disrupted using two cycles of 45 seconds at 30 Hz (Qiagen TissueLyser II). RNA was extracted with TRIzol Reagent (ThermoFisher Scientific) following the manufacturer’s procedure, treated with DNase (ThermoFisher Scientific), and quantified via the Qubit Fluorometric Quantification System High Sensitivity Assay (ThermoFisher Scientific). RNA concentration was standardized and then reverse transcribed using a High-Capacity cDNA Reverse Transcription Kit (ThermoFisher Scientific). Samples for Figure 7 were cleaned up with RNA Clean & Concentrator (Zymo) prior to quantification and cDNA synthesis.

### Quantitative RT-PCR assay and analysis

PCR primers were designed using Primer3web (http://www.bioinformatics.nl/cgi-bin/primer3plus/primer3plus.cgi; Untergasser et al, 2007) and Phytozome.com (https://phytozome.jgi.doe.gov/pz/portal.html) and ordered from Eurofins (https://eurofinsgenomics.com) (Supplemental Table 2). Quantitative PCR was carried out using SYBR Green MasterMix or PowerUP SYBR Green MasterMix (ThermoFisher Scientific) according to manufacturer’s instructions. Dissociation curves were analyzed to ensure that primers amplified a single product. Standard curves were generated using serial dilutions of pooled cDNA standards, data were analyzed using the relative standard curve method or ΔΔ*Ct* method and graphed with ggplot2 (v3.3.3) and ggpubr (v0.4.0) R packages.

### sgRNA design, cloning, and plasmid preparation

Coding sequences of *PpCDL1* and *PpCDL2* genes were used to search for CRISPR RNA (crRNA) preceded by a PAM motif of the *Streptococcus pyogenes* Cas9 (NGG or NAG) using the webtool CRISPOR V1 against the *P. patens* genome (http://crispor.tefor.net/). crRNAs within the first intron, with high specificity score, and few predicted off-targets, were selected for cloning (Supplemental Table 3). A fragment of 500 bp containing the snRNA U3 or U6 promoter (Collonnier et al. 2017) followed by a sgRNA, and flanked by AttB recombinant sites, was synthesized chemically as gBlocks (Integrated DNA Technologies) (Supplemental Table 4). sgRNAs are composed of 20 bp of the crRNA fused to 83 bp of the *S. pyogenes* tracrRNA scaffold (Mali et al. 2013). Each fragment was cloned into a pDONR207 (Invitrogen) backbone. Plasmids were amplified in *Escherichia coli* DH5a and purified using the NucleoSpin Plasmid Miniprep kit (Macherey-Nagel).

### Moss culture and transformation

Wild type *P. patens* Cholsey-K5, Trondheim-K2, and Reute accessions were used for transformation experiments. Plants were grown on solid BCD medium supplemented with 5 mM ammonium tartrate and were macerated multiple times to increase the amount of protonemal cells. Protoplast isolation and transformation followed (Roberts et al. 2011) with minor modifications. Protoplasts were transformed with 20 µg total circular DNA and regenerated following Collonnier et al. (2017) with minor modifications.

### DNA extraction, PCR screening, and sequencing of on-target mutations

Primers were designed to flank the sgRNA on-target sequence and to amplify ∼175 bp amplicon using Primer3web (http://www.bioinformatics.nl/cgi-bin/primer3plus/primer3plus.cgi) and Phytozome.com (https://phytozome.jgi.doe.gov/pz/portal.html). PCR of regions of interest was conducted with Plant Phire Direct PCR Master Mix (ThermoFisher Scientific) according to manufacturer’s instructions and evaluated via agarose gel electrophoresis. Products that differed in size from the wild type parent were sequenced via Sanger sequencing and analyzed using FinchTV v1.3.0 and AliView v1.25. Size-selective purification of amplicons was achieved by polyethylene glycol precipitation (Simpson 2006). Briefly, 50uL of 30% PEG-8000 30mM MgCl2 was added directly to PCR reaction. The total volume was adjusted to 100uL with TE. PCR amplicons were precipitated by centrifugation at high speed for 15 minutes. The DNA pellet was washed with 70% ethanol.

### Analysis of reproductive development of *Ppcdl* mutants

Transformants were transferred to 8L16D 15°C, and spores were collected for long-term storage in 500 microliters of sterile distilled water. Spore stocks were kept at 4°C in the dark. Spores from selected transformants and the wild type parent accession were plated on solid BCD medium at an empirically determined concentration such that approximately 10 spores germinated per technical replicate. These plants were grown in standard 16L8D 22°C conditions for 24 days, at which point replicates were vegetatively mature, before transfer to 12L12D at 22°C. Reproductive development was evaluated once per week for 24 weeks using a developmental scale (Supplemental Table 5), and data were graphed and analyzed using the ggplot2 (v3.3.3), ggthemes (v4.2.4), gridExtra (v2.3), and grid (v4.0.5) R packages.

### Yeast two-hybrid assay

Yeast two-hybrid tests were performed according to the Matchmaker® Gold Yeast Two-Hybrid System User Manual (Takara Bio Inc.). Briefly, *P. patens* prey coding sequences (CDL1, Pp3c3_26370; CDL2, Pp3c13_21190; LOVc15, Pp3c15_8610; FKc16, Pp3c16_19240; FKc25, Pp3c25_6710) were cloned into pGADT7-AD vectors. Similarly, these genes along with a *P. patens* Kelch coding sequence (Pp3c16_19240 excluding the F-box) were cloned into the bait vector, pGBKT7-BD. Plasmids were sequenced to confirm translational fusions. pGADT7 and pGBKT7 plasmids were transformed into yeast strains Y187 and Y2HGold, respectively, and yeast were mated according to Galletta and Rusan (2015) by interactions of interest. Diploid cultures were plated on -Leu/-Trp dropout media and -Leu/-Trp/-Ade/-His dropout media to select for yeast hybridization and protein interaction, respectively.

## Supporting information

Freidinger_et_al_supplemental

## Acknowledgements

We would like to thank Viktoria Coneva for her technical help while conducting experiments and advice on data analysis, Sofia Rehrig and Jackson Newell for assisting with data collection, Maria Sorkin for culturing tissue and preparing RNAs for the diurnal expression experiment, and members of the Kenyon College BIOL264 class for constructing several yeast two hybrid plasmids. This work was supported by NSF Grant 1656091, Kenyon Summer Science Scholars, and Kenyon College Department of Biology.

## Author Contributions

Karen A. Hicks, Lauren A. Woodward, and Alexander G. Freidinger conceived the project and designed the experiments. Alexander G. Freidinger, Lauren A. Woodward, Jo T. Bùi, Gillian Doty, Shawn Ruiz, Erika Conant, and Karen A. Hicks conducted the experiments and performed the data analysis. Alexander G. Freidinger, Lauren A. Woodward, Gillian Doty, and Karen A. Hicks wrote and edited the manuscript. All authors read and approved the manuscript.

