## Supplementary material for "Cycling DOF Factor mediated seasonal regulation of sexual reproduction and cold response is not conserved in *Physcomitrium patens*": Freidinger_et_al_supplemental

**Supplemental figures/tables**

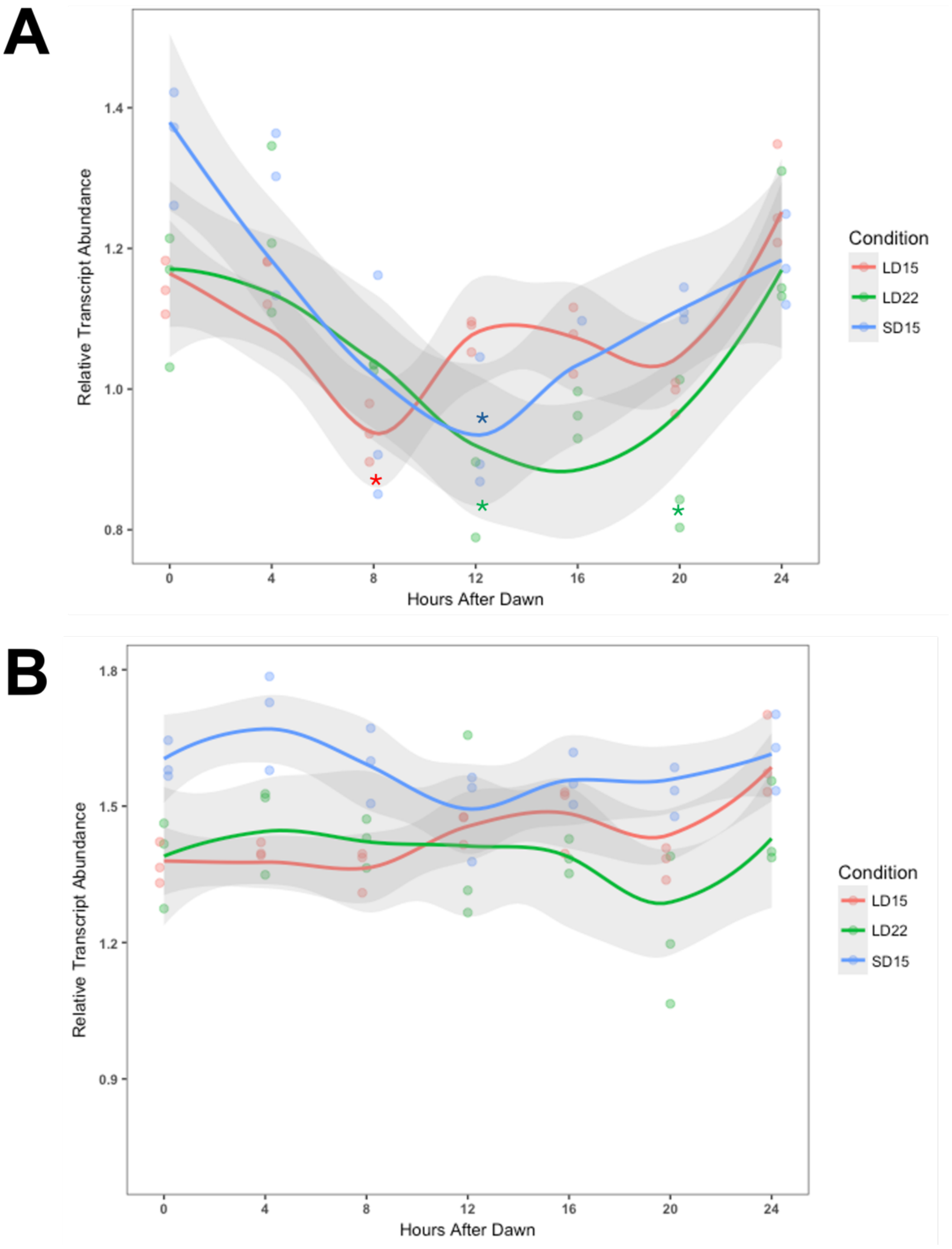


Supplemental Figure 1. *PpCDL1* is diurnally expressed in multiple environmental conditions while *PpCLD2* is not. A: Relative expression of *PpCDL1* is diurnal in 16L8D 22°C, 16L8D 15°C, and 8L16D 15°C in Villersexel-K4. Asterisks represent time points significantly different from ZT0 (ANOVA, p≤0.05). B: *PpCDL2* is not diurnally expressed in 16L8D 22°C, 16L8D 15°C, and 8L16D 15°C in Villersexel-K4. No time points were determined to be significantly different from ZT0 (ANOVA, p>0.05).

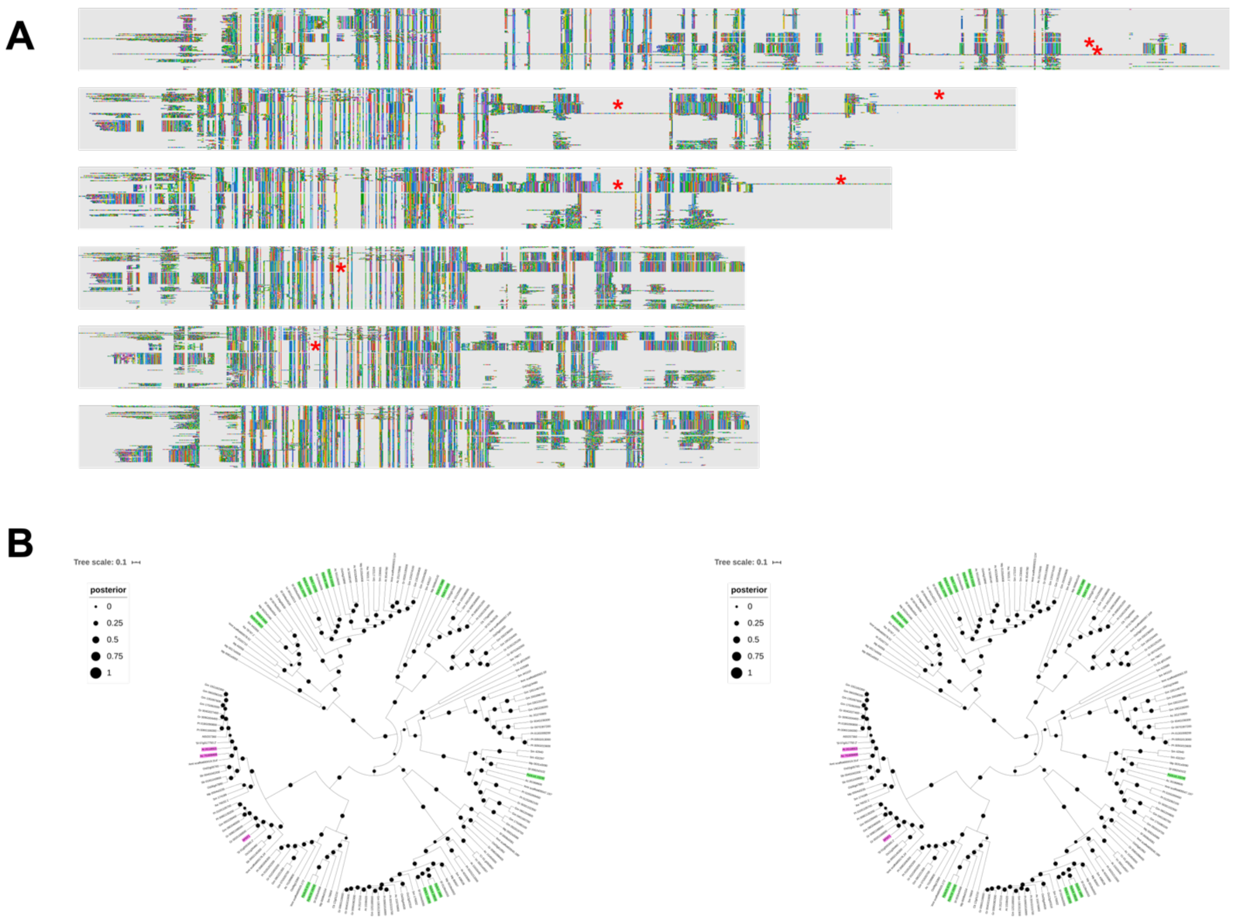


Supplemental Figure 2. Trimming protein alignments resulted in trees supported by higher posterior probabilities. A: Trimming of alignments used to generate the FKF1 phylogenetic tree shows larger continuous regions of amino acid conservation. Sequences with the only region of similarity being the DOF domain were removed. B: Comparison of trees generated with first and last alignments in panel A reveals increased confidence in tree topology. Left tree built using top alignment had posterior probabilities between 0.31 and 1.0. Right tree using bottom alignment had posterior probabilities between 0.51 and 1.0.

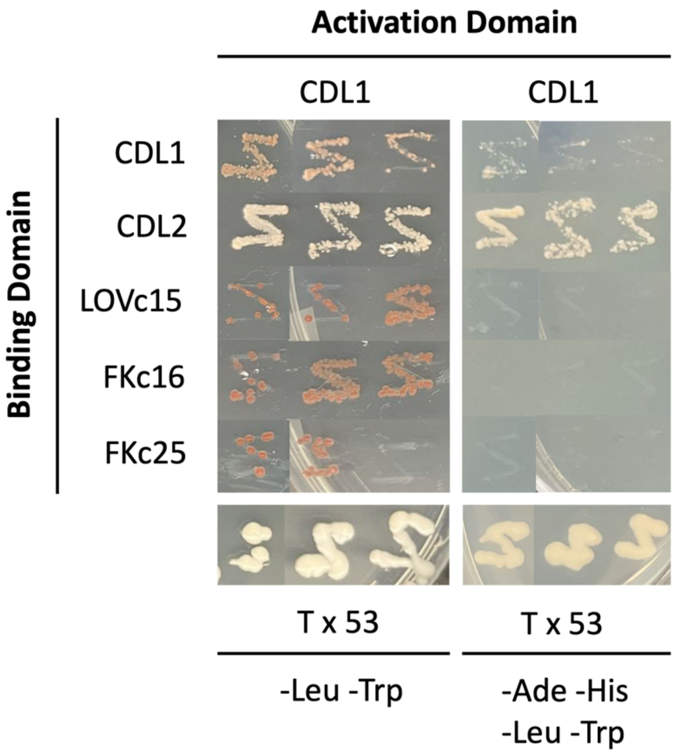


Supplemental Figure 3. A secondary plating of CDL1 x CDL2 growth from original Yeast two-hybrid assay confirms the interaction of these proteins in yeast. T x 53 is the positive control. -Leu/-Trp plates select for diploid yeast; -Ade/-His/-Leu/-Trp plates select for protein interaction.

Supplemental Table 1. Abbreviations for species used for phylogenetic analyses. Each sequence in phylogenetic figures is preceded by two or three letter species code.

| **Full Name** | **Abbreviation** |
|---|---|
| *Selaginella moellendorffii* | Sm |
| *Arabidopsis thaliana* | At |
| *Gossypium raimondii* | Gr |
| *Populus trichocarpa* | Pt |
| *Glycine max* | Gm |
| *Amborella trichopoda* | Amt |
| *Sphagnum fallax* | Sf |
| *Physcomitrella patens* | Pp |
| *Marchantia polymorpha* | Mp |
| *Aquilegia coerulea* | Ac |
| *Oryza sativa* | Os |
| *Solanum lycopersicum* | Sl |
| *Sorghum bicolor* | Sb |
| *Chlamydomonas reinhardtii* | Cr |
| *Volvox carteri* | Vc |
| *Chara braunii* | Cb |
| *Anthoceros agrestis* | Aa |

Supplemental Table 2. Primer sequences used for qRT-PCR analysis of *PpDOF* expression. Primers were designed using Primer3 and Phytozome. Primers that span an intron are noted with primer sequence-Intron-primer sequence.

| Gene | Sequence | Forward/Reverse |
|---|---|---|
| Pp3c3_26370 | ACCATGAGGATGAT-Intron-GGGAAA | Forward |
| Pp3c3_26370 | CCTTGGCCTTTGAATGAATG | Reverse |
| Pp3c13_21190 | GTTCGTCTCTG-Intron-GAACTTGCG | Forward |
| Pp3c13_21190 | TTCATTGTCCGCAACTGCA | Reverse |
| Pp3c3_950 | CATCCAAGCAG-Intron-GATGGAGGG | Forward |
| Pp3c3_950 | TCCCTGGAACCAGTTCAGTGG | Reverse |
| Pp3c4_850 | CTTGAATTGCAGAGGCAGTGAG | Forward |
| Pp3c4_850 | CCTTCAATCCAGCTCCCATTCC | Reverse |
| Pp3c5_14440 | GTGTACGGACAG-Intron-AGTCCGAATT | Forward |
| Pp3c5_14440 | CTTTCCATCATGAGCCATGCTATG | Reverse |
| Pp3c9_4910 | GGATCACGCAAGAAGCACAGTC | Forward |
| Pp3c9_4910 | AGGGTTGCCGATCTGCTCCAT | Reverse |
| Pp3c12_22390 | TTTGTGTGCGCTCGAGGTCAG | Forward |
| Pp3c12_22390 | TGGACTCGCCCACAGATCGATA | Reverse |
| Pp3c15_3180 | AGCGATGTCAAGGAGGCAAACC | Forward |
| Pp3c15_3180 | CAACTTGCTGCCGATGACGTAA | Reverse |
| Pp3c16_6460 | CCCATTAGAAAAG-Intron-GTATCTGCTACG | Forward |
| Pp3c16_6460 | CTGTTGAGATGTGCGACC | Reverse |
| Pp3c13_2000 | GCCTTCAG-Intron-CTGTAGAAGCGAG | Forward |
| Pp3c13_2000 | GAGTGTCCTGCGTTTGCC | Reverse |
| Pp3c17_3860 | ACCAGTCAGCCTCGCCACTA | Forward |
| Pp3c17_3860 | GGAACACTGCTGCTTTGAGGAA | Reverse |
| Pp3c17_7930 | CCTCTCAG-Intron-GCTGGTAGCGAA | Forward |
| Pp3c17_7930 | GCGAGCTTCTCAGAGGAGACTT | Reverse |
| Pp3c26_1490 | GCGTATCGCTG-Intron-ACTGGTAGT | Forward |
| Pp3c26_1490 | TTGACATCCTTAATCGCC | Reverse |

Supplemental Table 3. crRNA sequences designed using CRISPOR and selected to minimize off-site effects.

| ***P. patens* Gene** | **Strand** | **crRNA sequence** | **PAM Motif** | **Off-target Score** |
|---|---|---|---|---|
| *Pp3c3_26370* | Forward | GCTCTTCTGCAGACACCGGG | GGG | 0-0-0-0-2 |
| *Pp3c13_21190* | Reverse | GTGGCAGGCTTCGAAACCCT | AGG | 0-0-0-2-9 |

Supplemental Table 4. Sequences used in gene blocks for sgRNAs to create *ppcdl* mutants. Sequences were generated using CRISPOR V1 against *P. patens* genome v3. crRNAs within the first intron, with high specificity score, and few predicted off-targets, were inserted into fragment of 500 bp containing the snRNA U3 or U6 promoter (Collonnier et al. 2017) and flanked by AttB recombinant sites.

| Target gene | sgRNA sequence |
|---|---|
| *Pp3c3_26370* | ggggacaagtttgtacaaaaaagcaggcttcGAGCTCGAATTCGTCCATTGAAGCAGACGTGTTGCGA CAGGTTAGCGACGATGGGTGTAGATGTGATGTGATGTGATGGTGTGGTTCTTCCACG GCGGCGTCCTTGCGGTGGCGGAGAAGGGGATATCCCGAAGGAGCGGCAGCGGGAG AGCACAAGCAGAAAGGGTGCAGTGAGTGAGTGGGTCCAGCTGGGTGGCTGGCCGA GTGGACGCGACCGGGTTTCGAGGGGGcGGGGGAGAAAAGGGATGGAGCGAGGGAT ATAACCCACATGGAATGGAGGTGGGTGTGAAGGCGGGTATATAGGAAGGTGGAGGA CTTACAACCCATgCTCTTCTGCAGACACCGGGGTTTTAGAGCTAGAAATAGCAAGTT AAAATAAGGCTAGTCCGTTATCAACTTGAAAAAGTGGCACCGAGTCGGTGCTTTTTT TGAGCTCGTCgacccagctttcttgtacaaagtggtcccc |
| *Pp3c13_21190* | ggggacaagtttgtacaaaaaagcaggcttcGAGCTCGAATTCGTCCATTGAAGCAGACGTGTTGCGA CAGGTTAGCGACGATGGGTGTAGATGTGATGTGATGTGATGGTGTGGTTCTTCCACG GCGGCGTCCTTGCGGTGGCGGAGAAGGGGATATCCCGAAGGAGCGGCAGCGGGAG AGCACAAGCAGAAAGGGTGCAGTGAGTGAGTGGGTCCAGCTGGGTGGCTGGCCGA GTGGACGCGACCGGGTTTCGAGGGGGcGGGGGAGAAAAGGGATGGAGCGAGGGAT ATAACCCACATGGAATGGAGGTGGGTGTGAAGGCGGGTATATAGGAAGGTGGAGGA CTTACAACCCATgTGGCAGGCTTCGAAACCCTGTTTTAGAGCTAGAAATAGCAAGTT AAAATAAGGCTAGTCCGTTATCAACTTGAAAAAGTGGCACCGAGTCGGTGCTTTTTT TGAGCTCGTCgacccagctttcttgtacaaagtggtcccc |

Supplemental Table 5. Developmental maturity scale

| Phenotype | Description | Numeric score |
|---|---|---|
| 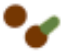 | Spores | 1 |
| 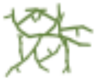 | Protonema | 2 |
| 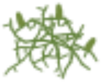 | Protonema with buds | 3 |
| 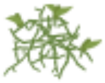 | Protonema with leaflets | 4 |
| 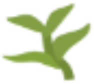 | Gametophores | 5 |
| 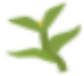 | Antheridia visible | 6 |
| 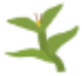 | Archegonia visible | 7 |
| 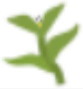 | Fertilized archegonia | 8 |
| 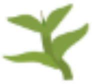 | Rounding arch | 9 |
| 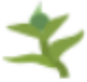 | Green sporophyte | 10 |
| 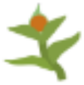 | Orange sporophyte | 11 |
| 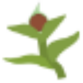 | Brown sporophyte | 12 |

Supplemental Table 6. Genotyping data of CRISPR-Cas9 modified Cholsey-K5 accession strains used in experiments. Each strain genotype was validated by Sanger sequencing of PCR-amplified genomic DNA using primers flanking the site of the expected mutation. Start codon (green), and guide RNA-targeted sequence (red) are shown in the genomic sequences of *PpCDL1* and *PpCLD2* (*Physcomitrium patens* genome v3.3) aligned with Sanger sequencing data from mutant strains.

<table>
<tr><th>Strain</th><th>cdl locus</th><th>Genotype</th><th>Sequencing alignment</th></tr>
<tr><td rowspan="2">double</td><td>ppcdl1</td><td>10bp deletion</td><td>M D S S S A D T G G D K P R V L K S A T<br>AATGGATAGCTCTTCTGCAGACACCGGGGGGGACAAGCCTAGGGTTTTGAAATCTGCCAC<br>||||||||||||||||||||||||          ||||||| ||||||||||||||||||<br>AATGGATAGCTCTTCTGCAGACAC----------AAGCCTAAGGTTTTGAAATCTGCCAC<br>M D S S S A D T S L R F * N L P L</td></tr>
<tr><td>ppcdl2</td><td>14bp deletion</td><td>N R M * T M G S S S H E N G G D E P R V<br>AACAGGATGTGAACAATGGGTAGCTCATCTCATGAAAACGGCGGTGACGAGCCTAGGGTT<br>||||||||||||||||||||||||||||||||||||||||||||||||||<br>AACAGGATGTGAACAATGGGTAGCTCATCTCATGAAAACGGCGGTGACGA----------<br>N R M * T M G S S S H E N G G D E<br><br>S K P A T V F L G D L S L S G E G P G F<br>TCGAAGCCTGCCACTGTTTTTTTGGGAGACTTGAGTTTAAGTGGTGAAGGGCCTGGATTT<br>    |||||||||||||||||||||||||||||||||||||||||||||  |||||||||<br>----AGCCTGCCACTGTTTTTTTGGGAGACTTGAGTTTAAGTGGTGAAGAACCTGGATTT<br>A C H C F F G R L E F K W * R T W I *</td></tr>
<tr><td rowspan="2">ppcdl2</td><td>ppcdl1</td><td>wt</td><td>M D S S S A D T G G D K P R V L K<br>AATGGATAGCTCTTCTGCAGACACCGGGGGGGACAAGCCTAGGGTTTTGAA<br>|||||||||||||||||||||||||||||||||||||||||||||||||||<br>AATGGATAGCTCTTCTGCAGACACCGGGGGGGACAAGCCTAGGGTTTTGAA<br>M D S S S A D T G G D K P R V L K</td></tr>
<tr><td>ppcdl2</td><td>13bp deletion</td><td>N R M * T M G S S S H E N G G D E P R V<br>AACAGGATGTGAACAATGGGTAGCTCATCTCATGAAAACGGCGGTGACGAGCCTAGGGTT<br>|||||||||||||||||||||||||||||||||||||||||||||||||||||||||<br>AACAGGATGTGAACAATGGGTAGCTCATCTCATGAAAACGGCGGTGACGAGCCTAGG---<br>N R M * T M G S S S H E N G G D E P R<br><br>S K P A T V F L G D L S L S G E G P G F<br>TCGAAGCCTGCCACTGTTTTTTTGGGAGACTTGAGTTTAAGTGGTGAAGGGCCTGGATTT<br>          ||||||||||||||||||||||||||||||||||||||||||||||||<br>----------CCACTGTTTTTTTGGGAGACTTGAGTTTAAGTGGTGAAGGGCCTGGAT--<br>P L F F W E T * V * V V K G L D</td></tr>
<tr><td rowspan="2">Transformed wt</td><td>ppcdl1</td><td>wt</td><td>M D S S S A D T G G D K<br>AATGGATAGCTCTTCTGCAGACACCGGGGGGGACAAG<br>|||||||||||||||||||||||||||||||||||||<br>AATGGATAGCTCTTCTGCAGACACCGGGGGGGACAAG<br>M D S S S A D T G G D K</td></tr>
<tr><td>ppcdl2</td><td>wt</td><td>AACAGGATGTGAACAATGGGTAGCTCATCTCATGAAAACGGCGGTGACGAGCCTAGGGTT<br>||||||||||||||||||||||||||||||||||||||||||||||||||||||||||||<br>AACAGGATGTGAACAATGGGTAGCTCATCTCATGAAAACGGCGGTGACGAGCCTAGGGTT<br><br>TCGAAGCCTGCCACTGTTTTTTTGGGAGACTTGAGTTTAAGTGGTGAAGGGCCTGGATTT<br>||||||||||||||||||||||||||||||||||||||||||||||||||||||||||||<br>TCGAAGCCTGCCACTGTTTTTTTGGGAGACTTGAGTTTAAGTGGTGAAGGGCCTGGATTT</td></tr>
</table>
